# Exon-based phylogenomics strengthens the phylogeny of Neotropical cichlids and identifies remaining conflicting clades (Cichliformes: Cichlidae: Cichlinae)

**DOI:** 10.1101/133512

**Authors:** Katriina L. Ilves, Dax Torti, Hernán Lépez-Fernández

## Abstract

The phenotypic, geographic, and species diversity of cichlid fishes have made them a group of great interest for studying evolutionary processes. Here we present a targeted-exon next-generation sequencing approach for investigating the evolutionary relationships of cichlid fishes (Cichlidae), with focus on the Neotropical subfamily Cichlinae using a set of 923 primarily single-copy exons designed through mining of the Nile tilapia (*Oreochromis niloticus*) genome. Sequence capture and assembly were robust, leading to a complete dataset of 415 exons for 139 species (147 terminals) that consisted of 128 Neotropical species, six African taxa, and five Indo-Malagasy cichlids. Gene and species trees were calculated using alternative partitioning schemes and reconstruction methods. In general, all methods yielded similar topologies to previously hypothesized relationships within the Cichlinae and clarified several relationships that were previously poorly supported or in conflict. Additional work will be needed to fully resolve all aspects of Cichlinae phylogeny. Overall, this approach yielded a well-resolved phylogeny of Neotropical cichlids that will be of utility for future assessments of the evolutionary and ecological processes within this diverse group of fishes. Furthermore, the general methodology employed here of exon targeting and capture should be applicable to any group of organisms with the availability of a reference genome.

## 1 Introduction

Neotropical cichlid fishes are rapidly becoming a model to understand the evolutionary history and biogeography of the exceptionally diverse Neotropical freshwater fish fauna (e.g., Arbour and López-Fernández 2014, 2016; Astudillo-Clavijo et al. 2015; Burress 2016; Hulsey and García de León 2005; Hulsey et al. 2006; López-Fernández et al. 2013; McMahan et al. 2013; Říčan et al. 2013; Tagliacollo et al. 2015). Likewise, the emergence of Neotropical cichlids as models of adaptive diversification in riverine environments is starting to provide a meaningful complement to the long-established studies of adaptive radiation of lacustrine cichlids in Africa (e.g., Brawand et al. 2014; Fryer and Iles 1972; Kocher 2004; Seehausen 2015; Wagner et al. 2012).

Nevertheless, continuing the evolutionary study of Neotropical cichlids depends on the availability of a robust phylogenetic framework that allows reliable reconstruction of divergence times, supports comparative analysis of lineage and phenotype divergence, and clarifies our understanding of biogeographic history.

Numerous studies have addressed the intergeneric and higher level relationships of Neotropical cichlids or some of their clades, and a relatively clear phylogenetic structure for the subfamily has emerged over the last two decades (e.g. Concheiro-Pérez et al. 2007; Farias et al. 1999, 2000; Hulsey et al. 2004, 2010; Kullander 1998; López-Fernández et al. 2010; McMahan et al. 2013; Musilová et al. 2008, 2009; Říčan et al. 2013, 2016; Smith et al. 2008). While these analyses have resulted in an increasingly stable understanding of Cichlinae relationships, a well-established taxonomy at the tribe level, and a relatively robust set of relationships among genera, a fully resolved and unambiguously supported phylogeny of Neotropical cichlids has yet to be achieved. This is particularly true of several basal relationships among genera or groups of genera within the three main tribes, Geophagini, Cichlasomatini and Heroini that remain poorly resolved or supported (e.g., López-Fernández et al. 2010; McMahan et al. 2013; Říčan et al. 2016).

Most analyses of Neotropical cichlid phylogeny have been based on relatively few loci (usually 10 or less) and often have been heavily informed by mitochondrial data (e.g., Friedman et al. 2013; López-Fernández et al. 2010; McMahan et al. 2013; Říčan et al. 2008; Smith et al. 2008). These studies are therefore limited in their ability to provide robust phylogenetic analyses, especially in the light of sequence saturation (e.g. Farias et al. 2001; López-Fernández et al. 2005), conflicting signal between nuclear and mitochondrial data (Dornburg et al. 2014; Říčan et al. 2016), and extensive basal short branches that often receive poor statistical support (López-Fernández et al. 2005, 2010). Beyond these well-known limitations, the small size of these datasets does not allow the incorporation of species tree approaches to phylogenetic analyses, and thus, all these studies are susceptible to producing misleading relationships due to conflict between gene trees and species trees (e.g. Edwards 2009; Maddison 2007). A recent study by Říčan et al. (2016) attempted to circumvent some of these potential limitations by using a large dataset of concatenated single nucleotide polymorphisms (SNPs) derived from restriction enzyme associated DNA (ddRAD). They obtained a largely well-supported tree with the largest taxon sampling of Central American cichlids to date, but their analysis was not able to unambiguously resolve some relationships and was limited to only one clade within Cichlinae. Moreover, Říčan et al.’s (2016) dataset is not amenable to be analyzed under coalescent-based methods and therefore cannot identify potential conflicts in phylogenetic relationships derived from the effects of deep coalescence (e.g. Edwards 2009; Heled and Drummond 2010). This latter point is relevant because the Neotropical cichlid phylogeny is plagued by short basal branches that, in previous work, have been interpreted as evidence of rapid diversification (López-Fernández et al. 2005, 2010). Concordant with this interpretation, comparative studies are revealing patterns of rapid early lineage and phenotype diversification through habitat and diet-related morphological diversification (e.g., Arbour and López-Fernández 2014; López-Fernández et al. 2013). These short branches, however, also represent a problem in phylogenetic reconstruction because rapid divergence can result in marked incongruence between gene divergence patterns and species divergence patterns due to population-level phenomena such as deep coalescence (e.g., Edwards 2009; Kubatko and Degnan 2007). Traditional phylogenetic methods such as those generally used in Neotropical cichlid analyses assume that loci in the genome diverge in concert with the species-level divergence of their corresponding species and that the phylogenetic signal is additive across loci; however, it is widely understood that genes have their own phylogenetic histories (gene trees) and that those histories do not always coincide node-to-node with the phylogenetic history of evolving lineages (species trees) (e.g., Maddison 2007). These incongruences between gene trees and species trees can be caused by several mechanisms, including hybridization and introgression, but for phylogenetic purposes the most widespread and potentially problematic is incomplete lineage sorting (ILS) or deep coalescence (Edwards 2009). From a phylogenetic reconstruction point of view, the challenge is how to identify the signal of individual gene divergence that is congruent with species divergence (Heled and Drummond 2010). It has been repeatedly shown that concatenation of sequences from genes with incongruent gene trees can produce erroneous topologies with high statistical support (Edwards 2009; Heled and Drummond 2010; Mendes and Hahn 2017). In the case of several Neotropical cichlid clades, short, poorly supported basal branches could result from poor sampling of that portion of the tree or represent times of divergence during which ILS resulted from fast species divergence. In the latter case, concatenated phylogenetic analyses could result in misleading topological reconstructions. Clearly separating these two scenarios, lack of data versus deep coalescence, may not be entirely possible, but the use of many independent nuclear loci (to increase gene tree sampling) and comparing concatenation methods and gene tree methods should help clarifying whether the pattern observed is due to a real evolutionary process or is the result of lack of data, improper analysis or both. Until recently, both the availability of methods to analyze species trees and the technical difficulties associated with identifying and sequencing many nuclear loci made this type of analysis very difficult or plainly inaccessible. With the advent of massively parallel sequencing and the development of methods that allow reconstructing phylogenies using models based on the coalescent, it is now possible to re-evaluate the phylogeny of Neotropical cichlids using phylogenomic datasets of hundreds of loci along with methods that correct for the effects of ILS. Despite these promising developments, however, computational limitations still restrict the options available to analyze truly phylogenomic datasets (e.g. hundreds or thousands of loci) with relatively large numbers of terminals and explicit coalescent analyses of sequence alignments.

In this study we used massively parallel sequencing and a custom-designed exon target-capture toolkit (Ilves and López-Fernández 2014) to generate a phylogenomic analysis of Neotropical cichlids under both concatenation and summary coalescent approaches. The main goals of the study were to update currently available hypotheses of Cichlinae relationships with a much larger dataset than previously available, and to leverage the phylogenomic dataset to perform coalescent-based analyses that should account for the possible effects of incomplete lineage sorting on our ability to accurately reconstruct Cichlinae relationships.

## 2 Materials and Methods

### 2.1 Taxon selection

Selection of taxa aimed at including as many lineages of Neotropical cichlids as possible. Representatives of all recognized genera were included for all tribes except the Heroini. A recent proliferation of new generic names for taxa within Heroini has taxonomically separated several monophyletic lineages that were previously considered congenerics (McMahan et al. 2015, Říčan et al. 2016). More importantly, many previously “orphan” clades variously referred to as ‘*Cichlasoma’ sensu lato* or *‘Heros’* have been formally assigned names. Given that these changes occurred after the dataset for this study was generated, representatives for some of these new genera are absent, but our dataset includes members of every clade within Heroini. Moreover, the study revises the identification of a few lineages of questionable identity in a previous analysis led by the senior author of this paper (López-Fernández et al. 2010, and see Říčan et al. 2013). Although those changes did not greatly affect the overall results of those analyses, they did incorporate additional uncertainty to the resolution of relationships within a clade of Heroini and, therefore, it is pertinent to correct them.

The dataset used herein closely resembles that of López-Fernández et al. (2010), and focuses on improving sampling of portions of the tree that were most problematic in that study in hopes of improving resolution and support for uncertain groupings such as those among the Cichlasomatini *Acaronia* and *Laetacara,* as well as some basal relationships among Geophagini and the heroin clade “amphilophines” sensu López-Fernández et al. (2010). Taxon sampling also was designed to test the depth of phylogenetic divergence that could be resolved using the targeted capture probes used in this study (and see Ilves and López-Fernández 2014). At the most recent variation end of the spectrum, we included two individuals of several species to test whether the dataset had enough phylogenetic signal to identify species as monophyletic. We also tested the ability of the dataset to resolve species-level relationships within genera by including a comparatively large number of species within the large *Crenicichla-Teleocichla* clade of Geophagini.

Altogether, the dataset included 147 terminals representing 139 species. Of these, 128 were Neotropical species and six were African taxa, which represent the sister group of Cichlinae. These taxa include *Heterochromis multidens,* the sister to all African cichlids, and both the reference sequence and a newly generated set of sequences for the Nile Tilapia, *Oreochromis niloticus,* the species from which the probes for target capture were developed (Ilves and López-Fernández 2014). Finally, we included five Indo-Malagasy cichlids which are sister to the African-Neotropical clade. The Sri Lankan cichlid *Etroplus suratensis* was used as the outgroup in all analyses. *Etroplus* and its sister genus, *Pseudetroplus,* are part of the Etroplinae subfamily which forms the sister group to the rest of the family (e.g., López-Fernández et al. 2010; Sparks 2008; Sparks and Smith 2004; Stiassny 1991). The complete list of species, museum catalog numbers, accession numbers (when available) and general locality data are given in Appendix A.

### 2.2. Library preparation for exon capture and sequencing

DNA extraction procedures followed those from Ilves and López-Fernández (2014). Exon capture of 923 targets and sequencing was performed at the Donnelly Sequencing Centre (DSC) at the University of Toronto (http://dsc.utoronto.ca/dsc/index.html) led by D. Torti, using the probes and general protocol described in Ilves and López-Fernández (2014). A double-hybridization procedure of probes to templates was performed, as this was found to significantly increase yield in previous work (Ilves and López-Fernández 2014). Details about the probe clean-up, library preparation, and exon capture procedure can be found in Appendix B. Paired-end sequencing was conducted on an Illumina HiSeq platform.

### 2.3. Automation of data processing and analysis

Multiple custom scripts were used to automate data processing and analysis from the initial step of read quality control to the creation of maximum likelihood phylogenetic gene trees. An overview file that describes each step can be found in Appendix C and subsequent appendices contain the specific script files used to perform each particular task or set of tasks.

### 2.4. Sequence quality control, read assembly and consensus sequence generation

The general procedures and parameters for the quality control of the sequencing reads, assembly of the reads into contigs, and generation of consensus sequences for each exon and species follow those from Ilves and López-Fernández (2014). A custom script was run that automated read quality control, contig assembly, and consensus sequence generation (Appendix D). Briefly, the standalone version of PRINSEQ (Schmieder and Edwards 2011) was used to retain only high-quality reads based on read length and base quality, bowtie2 version 2.1.0 (Langmead and Salzberg 2012) was used to map the contigs to a set of reference sequences from the Nile tilapia *(Oreochromis niloticus)* genome (Appendix E), and SAMtools version 0.1.19-44428cd (Li et al. 2009) was used to generate consensus sequences of the assembled contigs. Custom scripts were used to convert the FASTQ files to FASTA files for alignment (Appendix F) and convert all low quality base calls to “N” and trim terminal “N”s (Appendix G). FASTA sequences were imported into Geneious version 7.1.8 (http://www.geneious.com, Kearse et al. 2012), from which only sequences with a minimum of 100 bp were exported for subsequent use in alignment and phylogenetic analyses.

### 2.5. Sequence alignment and gene tree and species tree phylogenetic analyses

A custom script was used to combine all sequences for each exon into a single file (Appendix H). Sequence alignment for each exon was performed with muscle version 3.8.31 (Edgar 2004) using default parameters and was automated using a custom script (Appendix I). Each alignment was then manually inspected for quality and completeness in Geneious version 7.1.8 (http://www.geneious.com, Kearse et al. 2012). Only alignments that included a sequence of at least 100 bp for each taxon were retained for phylogenetic analyses. The 32 opsin exon alignments in the target kit were excluded from all analyses because they represent a family of genes for which duplication and pseudogenization events have been documented, complicating their use in phylogenetics due to paralogy (e.g., Bowmaker 2008; Weadick et al. 2012).

Coding and non-coding regions of each alignment were inferred from the annotated reference genome. Although the probes, as originally designed, were intended to target exon-coding regions, analyses of early alignments during the design of the protocol revealed non-open reading frames for some sequences (see Ilves and López-Fernández 2014). Additionally, subsequent iterations of annotation in the Nile tilapia genome revealed that some regions originally annotated as coding exonic regions actually corresponded to non-coding fragments. Coding and non-coding regions were identified and correspondingly separated for analyses in this paper. Bootstrapped gene trees (1000 replicates) for each exon alignment were generated using RAxML version 8.0.10 (Stamatakis 2014) with a GTRGAMMA model and a corresponding partition file with the codon positions and non-coding regions of the exon. This procedure was automated with a custom script (Appendix J).

ASTRAL-II version 4.8.0 (Mirarab and Warnow 2015) and STAR (Liu et al. 2009) species tree methods with multi-locus bootstrapping (Seo 2008) were run on the resulting bootstrapped RAxML datasets. ASTRAL-II analyses with 500 bootstrap replicates were run locally on a desktop Apple® iMac whereas STAR analyses were run on the STRAW server (Shaw et al. 2013). Because ASTRAL analyses require a maximum likelihood (ML) tree for each gene in addition to the set of bootstrapped tree files, RAxML was used to conduct 40 ML searches on each exon alignment (custom script Appendix K). To assess possible conflicts between species-tree and total-evidence concatenated analyses, we also performed an ML analysis of 415 concatenated loci with one thousand bootstrap replicates using RAxML.

### 2.6. Computing resources used

Read quality control, mapping, contig assembly, consensus sequence generation, and sequence alignment and some individual gene alignment RAxML analyses were performed locally on a desktop Apple iMac (3.5 GHz Intel Quad Core i7 with 32GB RAM and 3TB hard drive). Some individual gene alignment RAxML analyses were performed on the GPC supercomputer at the SciNet HPC Consortium (Loken et al. 2010).

### 2.7. Data availability

Alignments for all exons with complete taxon representation of 100 bp or greater as well as all bootstrapped phylogenetic trees, will be available on Dryad and all other data files, including raw fastq files, are available from corresponding author K.L. Ilves upon request.

## 3 Results

### 3.1. Target capture

923 exons are included in the probe set first developed by Ilves and López-Fernández (2014). 32 exons correspond to opsins, which were excluded from this study *a priori* due to their histories of gene duplication, which left a total of 891 exons as potential targets. Details about the number of reads obtained and retained after sequencing, the alignment rate, and number of exons captured per sample can be found in Appendix L (Table S1). Enforcing the restrictions of a complete dataset where every species must have a sequence of at least 100 bp, resulted in a dataset of 428 captured exons. 13 of these alignments were deemed to be ‘poor’ after visual inspection due to large blocks of ambiguous base pair calls (N) present in multiple species, often comprising over 50% of the total sequence length. Although the minimum sequence length was set at 100 bp, the average minimum sequence length relative to the average reference sequence length of 1136 bp was ∼62% (SD 23%), indicating that most species had sequence for most of the length of each target. The final dataset comprised 147 taxa representing 139 species for 415 exons (∼47% of target set) with a total length of 471,448 bp.

### 3.2. Depth of phylogenetic resolution in the exon-capture dataset

Analyses of multiple individuals within a species consistently resulted in monophyly at the species level with unambiguous support, as revealed by the concatenated analyses (Fig. S1, Appendix M). These results were consistent at various levels of divergence, ranging from resequencing of the African reference *Oreochromis niloticus* to newly sequenced Neotropical species in all major tribes (e.g. Geophagini: *Crenicichla sveni,* Cichlasomatini: *Krobia petitella,* Heroini: *Symphysodon aequifasciatus).* Both concatenated and consensus coalescence analyses of species-level divergence within genera also revealed a large phylogenetic signal that resulted in generally well-resolved and supported relationships among species within genera (e.g. *Crenicichla).* These results generally support the notion that the target capture probes used in this study are adequate to resolve phylogenetic relationships within Cichlidae spanning family to species levels of divergence.

### 3.3. Phylogenetic analyses

All analyses, regardless of methods, recovered the expected monophyletic Neotropical subfamily Cichlinae as sister to a monophyletic African Pseudocrenilabrinae, both of which in turn are sister to a paraphyletic arrangement of Indian and Malagasy lineages in the subfamilies Etroplinae and Ptychochrominae (Fig. 1). These relationships have long been well established through numerous studies of molecular and morphological datasets and will not be discussed further (Friedman et al. 2013; López-Fernández et al. 2010; McMahan et al. 2013; Matschiner et al. 2017; Smith et al. 2008; Sparks 2004, 2008; Stiassny 1991). Relationships among the seven recognized tribes of Neotropical cichlids (Cichlini, Retroculini, Astronotini, Chaetobranchini, Geophagini, Cichlasomatini and Heroini), were likewise recovered unambiguously across analyses. Results herein coincide with previous work in placing the Retroculini and Cichlini clade as sister to the remainder of Cichlinae. The tribes Cichlasomatini and Heroini form a monophyletic clade sister to a clade formed by Astronotini, Chaetobranchini and Geophagini. Interestingly, all analyses recovered Astronotini (genus *Astronotus)* as sister to the Chaetobranchini plus Geophagini clade. This relationship was also found by Matschiner et al. (2017), but has not been universally recovered in previous analyses: Smith et al. (2008) found *Astronotus* as sister to all Cichlinae except Cichlini and Retroculini, López-Fernández et al. (2010) found it as sister to the Cichlasomatini and Heroini clade, and McMahan et al. (2013) recovered the genus as sister to a clade of Chaetobranchini and Geophagini. All these studies were based on a limited set of genes and a large amount of mitochondrial data, suggesting that the much larger dataset used herein resolves the previously unstable placement of Astronotini.

**Figure 1.**
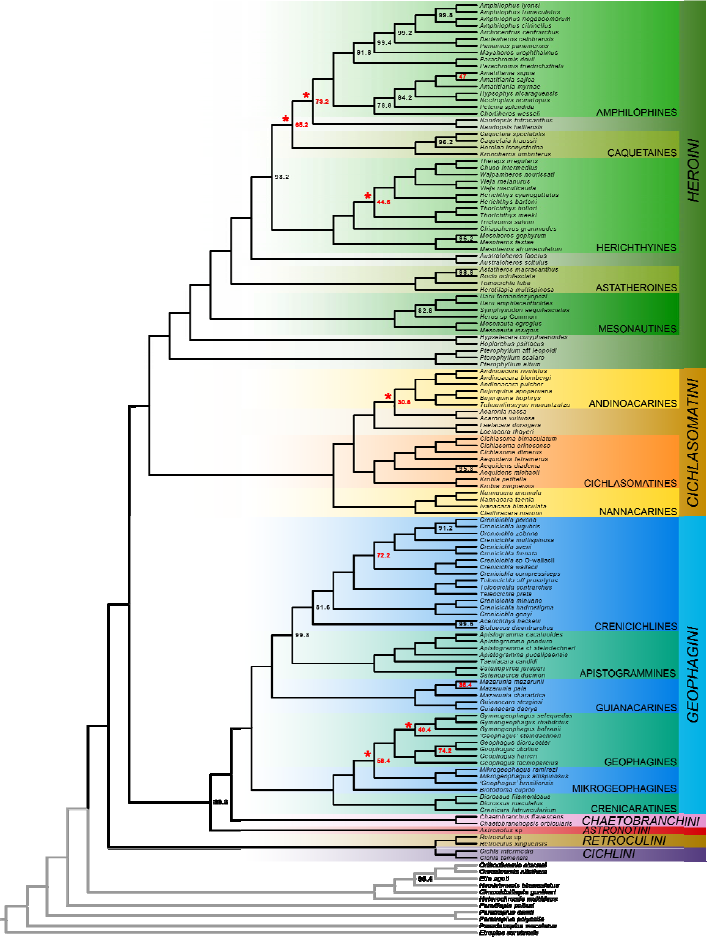
Species tree generated with ASTRAL-II for 415 loci comprising 471,448 bp. Colored clades depict formally recognized Neotropical cichlid tribes Heroini (Green), Cichlasomatini (Orange), Geophagini (Blue), Chaetobranchini (Magenta), Astronotini (Red), Retroculini (Brown), and Cichlini (Purple). See text for further discussion of relationships among and within tribes. Node bootstrap support is indicated when pertinent; nodes without labels received 100% support in this analysis. Nodes labeled in red received <75% bootstrap support. Nodes marked with an asterisk (*) represent weakly supported intergeneric relationships with incongruent resolution among two different species-tree and one concatenated phylogenetic analyses. See Figure 2 and Discussion for further analyses of these results. Complete topologies not shown here are provided in Figs. S1 and S2, Appendices M and N, along with node support.

In general, intergeneric relationships within each of the main Neotropical tribes, ‐‐ Geophagini, Cichlasomatini and Heroini‐‐, were recovered with unambiguous support. Nevertheless, considerable ambiguity was observed across methods in the placement of some lineages within these tribes (see Fig. 1). Given the widespread similarity among topologies, for the remainder of the paper we use the ASTRAL-II coalescent species-tree (Fig. 1) as a reference because of the superior performance of Astral-II as a tool for generating coalescent-based consensus species trees from gene trees (e.g. Arcila et al. 2016). Nevertheless, disagreements between this topology and those derived from the STAR species-tree method and the concatenated super-matrix topology are highlighted when pertinent (Fig. 2). Later we discuss the potential impact of topological uncertainty on macroevolutionary analyses and historical biogeographic studies of Neotropical cichlids.

**Figure 2.**
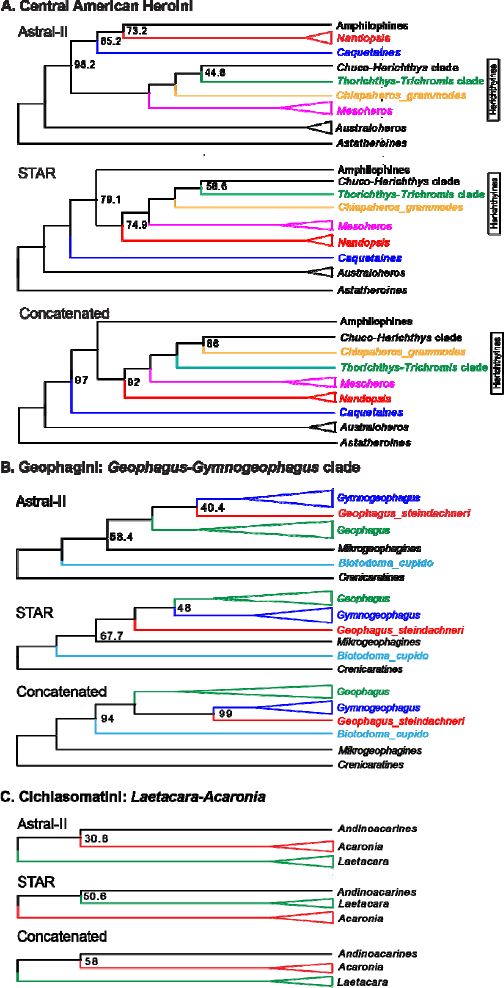
Conflicting results among analyses. Each panel depicts the alternative topological arrangements found in the clades highlighted with asterisks in Fig. 1. Colors are meant to represent lineages within each panel and not to be used as comparison among clades in different panels. Numbers by nodes represent bootstrap support in each case; nodes without numbers received 100% bootstrap support in their respective analyses. See Fig. 1 for the ASTRAL-II species tree and Appendices M and N (Figs. S1 and S2) to see the concatenated and STAR topologies, respectively.

Within the tribe Heroini, all analyses coincide in placing the South American lineages as a paraphyletic arrangement at the base of the clade, with the genus *Pterophyllum* as the earliest diverging lineage, followed by a clade of *Hoplarchus* and *Hypselecara* and then by a clade (mesonautines, Fig. 1) in which *Mesonauta* is sister to *Heros,* which is in turn sister to *Symphysodon* and *Uaru.* A similar arrangement has been found by other studies (e.g., McMahan et al. 2013), but the position of *Pterophyllum* can be flipped with that of *Hypselecara* and *Hoplarchus* clade (e.g., López-Fernández et al. 2010). Likewise, all analyses resulted in less than complete support for the position of *Heros,* but the placement does not vary across trees in this study. Support for the placement of *Heros* in previous studies (e.g., López-Fernández et al. 2010) was relatively weak, and despite orders of magnitude increase in the number of loci used herein, the phylogenomic analyses still result in less than 100% bootstrap support for the position of the genus. Contrastingly, the current analyses removed all ambiguity from the previously uncertain placement of the genus *Uaru* (López-Fernández et al. 2010).

As in previous studies, the remainder of Heroini is comprised by a geographically non-monophyletic arrangement of South and Central American lineages. Among these, a clade including the genera *Rocio, Tomocichla, Herotilapia* and *Astatheros* is sister to the rest of Heroini. This arrangement is equivalent to and in the same position of the astatheroines clade sensu Říčan et al. (2016), but it does not include their newly described genus *Cribroheros,* which was not represented in our dataset. Říčan et al. (2016) recently split *Astatheros* sensu López-Fernández et al.’s (2010) into *Astatheros, Rocio* and *Cribroheros,* but phylogenetically there is no incongruence among the groups in both studies. The astatheroine clade is sequentially followed by the genus *Australoheros* as sister to all other heroins, an arrangement identical with that found by Říčan et al’s (2016) analysis based on concatenated single nucleotide polymorphisms from a restriction site associated DNA (RAD) dataset. This result differs from that of López-Fernández et al. 2010 because the only species of *Australoheros* in that study was inadvertently switched with that of the unrelated *Cryptoheros nanoluteus* (see Appendix A for details, and Říčan et al. 2013). McMahan et al. (2013) also find a different placement for *Australoheros*.

Our analyses unambiguously find a monophyletic clade of purely Central American taxa that corresponds with the amphilophines of Říčan et al. (2016), even though our analysis did not include their genera *Cryptoheros, Talamancaheros* and *Isthmoheros.* Despite this correspondence, however, our analyses produced different internal relationships among the included genera. For example, while Říčan et al. (2016) found *Parachromis* in a subclade with *Amatitlania,* we find it in a different subclade that includes *Amphilophus* (Fig. 1). Likewise, our grouping corresponds roughly with the amphilophines of López-Fernández et al. 2010, but excludes *Trichromis (‘Cichlasoma’ salvini* in their Fig. 1). Even though the results of our phylogenomic analyses have stronger statistical support that those of López-Fernández et al. (2010) and of Říčan et al. (2016, see their Fig. 5), the amphilophine clade found herein contains the largest number of weakly supported nodes in our analyses. It is interesting to note that, among amphilophines, the sequential position of *Petenia* and *Chortiheros* as sister to a clade of *Amatitlania, Hypsophrys* and *Neetroplus* was consistently recovered in all analyses, but support for this arrangement was ambiguous (Figs. 1, S1 [Appendix M], and S2 [Appendix N]). Říčan et al. (2016) recovered *Petenia* and *Chortiheros* as sister to each other, but none of our analyses supported that relationship.

The largest disagreement among topologies obtained in this study, as well as with those from previously published analyses, involves the genera in the informal clade herichthyines, the Caribbean genus *Nandopsis,* and the caquetaines clade containing the genera *Caquetaia, Heroina* and the recently named genus *Kronoheros* (Fig. 2A). The relationship between these lineages and the amphilophines also varies among our three analyses, and often differs from relationships found in other studies. It is interesting that our concatenated and STAR coalescent analyses are more similar with each other than either is to the ASTRAL-II topology. In Říčan et al.’s (2016) analysis, the caquetaines were sister to a clade of amphilophines and herichthyines with *Nandopsis* sister to the latter. This is a similar arrangement to that found in our STAR and concatenated analyses, but in our ASTRAL-II topology (Fig. 2A) *Nandopsis* was sister to amphilophines, and in turn, the two were sister to the caquetaines. In all analyses, at least some of these relationships are inconclusively supported, although the concatenated analysis obtained the highest bootstrap values of the three. In Říčan et al’s (2016) analysis, *Chiapaheros* remained in a polytomy. In our analysis, it is part of a well-supported clade along with two other well-supported groupings: *Thorichthys* and *Trichromis* and a clade comprising *Herichthys, Vieja, Wajpamheros, Chuco* and *Theraps,* but the relationships among these three lineages remain unclear (Fig. 2A).

Relationships among Geophagini in all analyses were generally compatible with those previously described by López-Fernández et al. (2012) and comprised by two major clades compatible with those described by López-Fernández et al. (2010, 2012). In the first of these, a clade of *Guianacara* and *Mazarunia* is sister to a clade of “apistogrammines” and “crenicichlines” sensu López-Fernández et al. (2010). Except for a few nodes within the *Crenicichla-Teleocichla* group and another one within *Mazarunia,* relationships within this large clade are largely congruent with previous results and well supported. Additionally, the sister relationship between *Acarichthys* and *Biotoecus* was unambiguously recovered, but support of its sister relationship to *Crenicichla* was always below 100%.

A second clade of Geophagini comprised the genus *Biotodoma,* the “geophagines”, “mikrogeophagines” and “crenicaratines” of López-Fernández et al. (2010). Monophyly of the “geophagines” (genera *Geophagus, Gymnogeophagus* and the *‘Geophagus’ steindachneri* group) was unambiguously supported by all analyses, but relationships within the group were different in the three topologies (Fig. 2B). Moreover, the relative position of the genus *Biotodoma* and the “mikrogeophagines” (genera *Mikrogeophagus* and *‘Geophagus’ brasiliensis)* was different in the coalescent-based analyses compared to the concatenated topology. As discussed below, ambiguity in the placement of the “geophagines” genera could have implications for estimating the age of cichlids.

Relationships within Cichlasomatini were generally identical among analyses. In all cases, a clade including *Nannacara, Ivanacara* and *Cleithracara* was recovered as the unambiguously supported sister group to the rest of Cichlasomatini. This clade is equivalent to the “nannacarines” sensu López-Fernández et al. (2010), but its position with respect to the rest of the tribe is novel with respect to previous studies. Musilová et al. (2009) found the “nannacarines” as sister to *Laetacara* but with low support, and López-Fernández et al. (2010) found it as sister to a clade of *Laetacara, Acaronia* and their “andinoacarines”, but also with low support. The genera *Cichlasoma* and *Aequidens* were found as sister to *Krobia* in all studies and as expected from previous work (López-Fernández et al. 2010; Musilová et al. 2009). Our analyses also unambiguously recovered a well-supported monophyly of *Andinoacara* and *Bujurquina* which in turn are sister to *Tahuantinsuyoa* (andinoacarines, sensu López-Fernández et al. 2010). *Acaronia* and *Laetacara* were recovered as sister to the “andinoacarines” but the relative placement of the genera with respect to each other was not unambiguously supported in any of the analyses. Placement of these two genera has varied across studies: Musilová et al. (2009) found *Acaronia* to be sister to all Cichlasomatini, with *Laetacara* either sister to the remainder of the tribe or to the *Cichlasoma, Aequidens* and *Krobia* clade (cichlasomatines); López-Fernández et al. (2010) recovered a weakly supported sister clade that in turn was sister to “andinoacarines”, but support was low.

## 4. Discussion

Even though higher-level relationships among clades and genera of Neotropical cichlids have become increasingly resolved and supported by recent work, the position of several groups remains uncertain or weakly supported. In this study, we used a recently developed set of exon-targeting probes and massive parallel next generation sequencing to generate a large dataset aimed at resolving Cichlinae relationships that remain poorly supported. Our phylogenomic analyses confirm many relationships previously found among Neotropical cichlids, and provide unprecedented resolution and support for many relationships that were previously weakly supported, especially near the base of the tree. This is particularly true among some genera of Central American lineages in the amphilophines clade, the unambiguous clarification of the relationship among Geophagini clades crenicichlines, apistogrammines and guianacarines (sensu López-Fernández et al. 2010), and the strongly supported position of the Cichlasomatini clade nannacarines as the sister group to the rest of the tribe (Fig. 1 and see Figs. S1 and S2, Appendices M and N, respectively). Nevertheless, despite an increase of two orders of magnitude in data when compared with other sequencing studies (e.g., López-Fernández et al. 2010; Říčan et al. 2013), some relationships remain unclearly established or poorly supported. The three main regions of the tree that continue to resist clear resolution include the Central American herichthyines, the position of the Cichlasomatini genera *Laetacara* and *Acaronia,* and the order of divergence among genera in the Geophagini clade including *Geophagus, Gymnogeophagus* and the *‘Geophagus’ steindachneri* clade (Fig. 2).

Previous studies have repeatedly found that Neotropical cichlids diversified over a relatively short time period, as evidenced by short branches and frequently weakly supported relationships near the base of the tree (e.g., Farias et al. 1999; López-Fernández et al. 2005, 2010). Most of the previously unresolved or weakly supported relationships in the López-Fernández et al. (2010) study, which has the most comparable taxon sampling to this study, are resolved with unambiguous support by the phylogenomic analyses presented here regardless of the method used (compare Fig. 1 of both studies). Thus, in combination, the generalized stability of higher-level phylogenetic hypotheses obtained through both summary coalescent and concatenation methods in our analyses, suggests that conflict between gene trees and the species tree is relatively rare among Neotropical cichlids. It appears that incomplete lineage sorting and other sources of misleading phylogenetic signal, such as introgression or hybridization, do not frequently disrupt our ability to reconstruct relationships. We make this assertion with caution, however, because detailed phylogeographic studies of Neotropical cichlids have shown that at least some taxa may be affected by these problems, particularly at more recent levels of divergence (e.g., Willis 2017; Willis et al. 2013). Even more germane to our study, the persistence of unresolved or poorly supported “deep” relationships within a few clades of Cichlinae may indicate a role for deep coalescence or other confounding effects in some early events of divergence among Neotropical cichlids (Fig. 2).

This point is particularly relevant because the tribes containing the remaining unresolved or conflictive clades underwent relatively quick adaptive diversification giving origin to a variety of lineages (Arbour and López-Fernández 2014; López-Fernández et al. 2013). It is suggestive that the largest remaining conflicts within the tree involve the Central American herichthyines, including some Central American genera with South American distribution and the Caribbean genus *Nandopsis* (Fig. 2A). Recent work has shown that invasion of Central America by the tribe Heroini provided renewed ecological opportunity that allowed this clade to rapidly diversify into a broad variety of ecologically specialized forms (Arbour and López-Fernández 2016). It is conceivable that such rapid adaptive divergence produced incomplete lineage sorting among some heroine lineages, resulting in reduced phylogenetic resolution (e.g., Edwards 2009; Kubatko and Degnan 2007). In fact, the early radiation of the Neotropical cichlid tribes in South America has been similarly shown to have occurred quickly, potentially leaving a similarly conflictive gene tree divergence patterns in other regions of the tree, particularly within Geophagini, which dominates the lineage and functional diversity of South American Neotropical cichlids (Arbour and López-Fernández 2014; Astudillo-Clavijo et al. 2015; López-Fernández et al. 2013). Even if early adaptive radiation in Neotropical cichlids resulted in incomplete lineage sorting in some clades, our results suggest that its effects may not be extensive because, with the exceptions pointed out above, both our concatenated and coalescent analyses recover largely congruent phylogenies. Moreover, most of the relationships recovered herein are congruent with those founds in previous studies based on much smaller, concatenated datasets (e.g., López-Fernández et al. 2010; McMahan et al. 2013; Musilová et al. 2009; Říčan et al. 2008) and with the ddRAD-based analysis of Central American heroines of Říčan et al. (2016).

The ability to generate a reliable phylogeny has important consequences beyond the mere systematic implications of the study. Uncertainty about the order of divergence and relationships among genera and higher clades can affect our ability to reconstruct the history of evolutionary divergence in cichlids. Three types of studies could be affected by diminishing but still present uncertainty in Neotropical cichlid relationships. Robust topologies are critical for using the fossil record to calibrate molecular phylogenies and reconstruct the timeline of diversification of a clade. The fossil record of cichlids is relatively scarce, and considerable debate has ensued regarding the identity and placement of fossils, particularly the recently described Eocene fossils from the Lumbrera formation in Argentina (e.g., Friedman et al. 2013; López-Fernández et al. 2013; Malabarba et al. 2010, 2014). On the one hand, the strong support received by most nodes in our analyses should provide a solid scaffold for time calibration. Unfortunately, one of the most unstable relationships, that involving the geophagine genera *Gymnogeophagus, Geophagus* and *‘Geophagus’ steindachneri,* affects the certainty with which the Eocene fossil † *Gymnogeophagus eocenicus* can be placed on the tree. As a consequence, calibration of *Gymnogeophagus* remains uncertain because it is unclear whether the genus is sister to the broadly distributed *Geophagus* sensus stricto or to the northern Andes clade including *‘Geophagus’ steindachneri* (Fig. 2B). Therefore, the inconclusively established position of a fossil-bearing clade can have important consequences in both reconstruction of the timeline of divergence and the historical biogeography of South American cichlids.

From a biogeographic point of view, the uncertainty observed among Central American cichlids is likely to have even more dramatic consequences. The incongruent placement of the Caribbean genus *Nandopsis* and of the South American caquetaines limits our ability to accurately recreate the history of heroine invasion of Central America and for understanding the events driving the potential recolonization of South America by some of the Mesoamerican heroin lineages such as *Mesoheros* and the caquetaines. Recent studies that have addressed the historical biogeography of Central American cichlids and their relationships to South America have relied on single reconstructions of the phylogeny based on either a small number of concatenated loci (e.g., Říčan et al. 2013; Tagliacollo et al. 2015) or on phylogenomic approaches based on concatenation of single nucleotide polymorphisms (Říčan et al. 2016). These studies focus on interpretations of the particular trees found in each of their analyses, but our study suggest that the historical biogeography of Central America may require either a more exhaustive analysis of phylogenetic relationships or the reconstruction of competing, alternative scenarios that reflect the current uncertainty in the phylogeny. Based purely on methodological arguments of performance and accuracy (e.g., Arcila et al. 2017; Mirarab and Warnow 2015), it is possible that our ASTRAL-II topology provides a more stable framework for analysis, but the weak support received by nodes in that arrangement suggest that any interpretations of historical biogeography should be done cautiously.

Phylogenies are also becoming increasingly important as the framework to perform macroevolutionary analyses of lineage and phenotypic divergence. With a proliferation of comparative methods and the increased availability of well supported and increasingly better dated trees, our ability to infer patterns and processes of divergence continues to improve. With several studies recently addressing the evolution of Neotropical cichlids (e.g.,, Arbour and López-Fernández 2013, 2014, 2016; Astudillo-Clavijo et al. 2015; Burress 2016; Hulsey et al. 2006; López-Fernández et al. 2013), it is pertinent to ask whether changes in the topology may require modification of our emerging understanding of Cichlinae macroevolution. The phylogeny obtained herein is remarkably similar to the López-Fernández et al. (2010) tree used in most of the macroevolutionary analyses listed above, suggesting that macroevolutionary conclusions to date are robust. When analyses are performed on a sample of the posterior distribution of the dated phylogeny and not just on the maximum clade credibility (MCC) tree, topological uncertainty should be reflected in the results such that observed macroevolutionary patterns should be robust to moderate topological changes. This should be particularly true for analyses of phenotypic divergence based on the construction of morphospaces, such as disparity through time and adaptive landscape inferences (e.g., Ingram and Mahler 2013; Slater and Pennel 2014). Likely more sensitive to changes in the tree are lineage through time and rate analyses because they depend on branch lengths in ultrametric trees, which in turn depend on the accuracy of both the topology and of age estimates. Because coalescent summary analyses such as ASTRAL-II and STAR do not provide branch lengths, the species trees estimated here cannot be used directly in comparative analyses that require ultrametric topologies derived from estimates of absolute time.

Finally, generating an ultrametric topology by dating phylogenomic datasets remains a challenge. In principle, there is no reason that phylogenomic species-tree hypotheses cannot be dated, but in practice it is not clear if the incorporation of large phylogenomic datasets and fossil data into actual analyses is computationally feasible (e.g., Bouckaert et al. 2014; Matschiner et al. 2017). Moreover, among other problems, traditional node dating methods require a priori placement of fossils on topologies, further complicating the use of fossils in unresolved clades, such as *†Gymnogeophagus eocenicus* within the geophagines. Alternatively, emerging total evidence dating methods that simultaneously generate phylogenies and age estimates combining molecular, morphological and fossil data may provide more flexibility and accuracy *vis a vis* uncertainty in the molecular phylogenies (e.g., Heath et al. 2014). Whether these methods can be employed in truly phylogenomic contexts with hundreds of loci and taxa is not yet clear (e.g., Gavryushkina et al. 2016; Ronquist et al. 2012).

## 5. Conclusion

Phylogenomic analyses of Neotropical cichlids (subfamily Cichlinae) using 415 exons for 139 species in both concatenated and summary statistic coalescent frameworks resulted in generally well-resolved, strongly-supported and broadly-congruent topologies. The topologies obtained are similar to previous hypotheses of relationships among Cichlinae but, in general, provide stronger support for many relationships that previously had weak or conflicting support. The results of our analyses also suggest that the targeted genomic regions contain phylogenetic signal capable of resolving relationships at all levels of divergence within the clade. Nevertheless, we identified several regions of the tree in which relatively early divergence events cannot be reconstructed with certainty because different methods provide conflicting results, none of them with conclusive support. We suggest this incongruence may result from incompletely lineage sorting associated with the early adaptive divergence of the clades in which incongruence is observed. We argue that, despite some disagreements in the placement of these lineages both in this study and in previous analyses, most studies provide a broadly common signal of evolutionary relationships among Neotropical cichlids. A generally better supported phylogeny derived from our phylogenomic analyses should continue to provide a solid framework for the evolutionary analysis of lineage and phenotypic divergence in Neotropical cichlids. Future work will focus in further clarifying relationships within the few recalcitrant clades identified herein, and in leveraging the current dataset as a uniquely strong molecular framework for clarifying the timeline of evolutionary divergence among Neotropical cichlids.

## Acknowledgements

The authors are grateful to the following for curatorial assistance and the loan of tissue samples not available at the ROM collection: M. Burridge, M. Zur, and E. Holm (Royal Ontario Musem [ROM]), M. Stiassny and B. Brown (American Museum of Natural History, New York), M. Sabaj (Academy of Natural Sciences of Drexel University, Philadelphia), R. Rodíles-Hernández (ECOSUR), L.R. Malabarba (Universidade Federal do Rio Grande do Sul, Porro Alegre, Brazil), J. Armbruster (Auburn University Museum, Auburn, Alabama) and Yves Fermon (Muséum National d’Histoire Naturelle, Paris). We further thank the people who assisted during fieldwork related to this work: J. Arbour, D. Bloom, R. Rodíles-Hernández, F. Hauser, N. Lujan, N. Meliciano, E. Liverpool, C. Montana, G. Ortí, S. Refvik, M. Röepke, M. Soria, D. Taphorn, M. Tobler, S. Willis, and K. Winemiller. Some computations were performed on the GPC supercomputer at the SciNet HPC Consortium. SciNet is funded by the Canada Foundation for Innovation under the auspices of Compute Canada, the Government of Ontario, Ontario Research Fund - Research Excellence, and the University of Toronto. We further thank R. Williamson for assistance with the development in the custom scripts used in the bioinformatics pipeline. Funding for this work, including fieldwork and laboratory analyses was provided by a Discovery Grant from the Natural Sciences and Engineering Research Council of Canada, ROM Governors and the National Geographic Society grants to HLF. Additional fieldwork was supported by a Coypu Foundation Grant to N. Lujan and ROM Trust Funds to HLF. KLI was supported by a Rebanks Postdoctoral Fellowship from the ROM Governors granted to HLF to develop a phylogenomic toolkit for the phylogenetic analysis of cichlids.

